# Development and Binocular Matching of Orientation Selectivity in Visual Cortex: A Computational Model

**DOI:** 10.1101/682211

**Authors:** Xize Xu, Jianhua Cang, Hermann Riecke

## Abstract

In mouse visual cortex, right after eye-opening binocular cells have different orientation preferences for input from the two eyes. With normal visual experience during a critical period, these orientation preferences shift and eventually become well matched. To gain insight into the matching process, we developed a computational model of a cortical cell receiving - via plastic synapses - orientation selective inputs that are individually monocular. The model captures the experimentally observed matching of the orientation preferences, the dependence of matching on ocular dominance of the cell, and the relationship between the degree of matching and the resulting monocular orientation selectivity. Moreover, our model puts forward testable predictions: i) the matching speed increases with initial ocular dominance and decreases with initial orientation selectivity; ii) matching proceeds faster than the sharpening of the orientation selectivity, suggesting that orientation selectivity is not a driving force for the matching process; iii) there are two main routes to matching: the preferred orientations either drift towards each other or one of the orientations switches suddenly. The latter occurs for cells with large initial mismatch and can render the cell monocular. We expect that these results provide insight more generally into the development of neuronal systems that integrate inputs from multiple sources, including different sensory modalities.

**New & Noteworthy:** Animals gather information through multiple modalities (vision, audition, touch, etc). These information streams have to be merged coherently to provide a meaningful representation of the world. Thus, for neurons in visual cortex V1 the orientation selectivities for inputs from the two eyes have to match to enable binocular vision. We analyze the postnatal process underlying this matching using computational modeling. It captures recent experimental results and reveals interdependence between matching, ocular dominance, and orientation selectivity.

## 1 Introduction

Animals receive information about the world through multiple modalities (vision, audition, touch, etc). For these information streams to provide a meaningful representation of the sensory world they have to be merged in a coherent fashion; only then do they enable the brain to better detect events, analyze the corresponding situations and then make decisions accordingly. Typically, this coherence is only acquired during a postnatal critical period [1].

The merging of across-modality information has been extensively investigated in the cat superior colliculus (SC) [2, 3, 4, 5] as well as the optic tectum of the barn owl [6], where multisensory neurons integrate the information they receive from upstream unisensory neurons in different sensory channels (e.g., visual and auditory). Like in other sensory systems, the capability of SC multisensory neurons to engage in multisensory integration is not innate but learned gradually during postnatal life as a consequence of normal multisensory experience. Two main results of the multisensory neurons’ learning process in SC are the initial development of large, unisensory receptive fields for visual and auditory input and their subsequent contraction and matching across modalities. This can enhance the degree to which the neurons’ receptive fields for visual and auditory inputs pertain to the same spatial location and enables the neuron to extract coherent information from the different modalities.

Matching of different information streams can also play an important role within a single modality. In the visual system, for instance, neurons in the visual cortex prefer similar orientations through the two eyes. As in the multisensory case, this binocular matching requires normal sensory experience. Shortly after eye-opening cortical cells in layer 2/3 have quite different monocular orientation preferences through each eye [7]. With normal binocular visual experience these preferences become binocularly matched to the adult level by postnatal day 31 (P31) [7], which corresponds to the end of the critical period for ocular dominance plasticity [8].

Inspired by these experimental results and to gain insight into general matching mechanisms, we developed a computational model for the development and matching of input preferences of neurons receiving multi-channel input via plastic synapses. In the case of multisensory SC neurons the input preferences would correspond to visual and auditory receptive fields. For concreteness, we will focus here on the binocular matching in V1, where the input preferences correspond to orientation preferences.

We considered a single spiking neuron that receives orientation-selective inputs, separately from each eye. The evolution of the synaptic weights was driven by stimuli representing gratings with randomly switching orientation. In an initial phase these inputs were uncorrelated between the two eyes to mimic spontaneous retinal or thalamic activity before eye opening [9]. After eye-opening the inputs were chosen to be perfectly correlated between left and right. Our aim was to keep the model as simple as possible, while still capturing a wide range of experimental observations. We therefore did not modify the plasticity rules when switching between these two phases and did not include a transition period (P15-P20) during which the input changes from being dominated by spontaneous activity to being dominated by visually-evoked activity [10].

Our model captures key experimental observations [11, 12]:

1. the matching is predominantly achieved by shifting the orientation preference for input from the weaker eye.
2. the resulting binocular orientation selectivity increases with decreasing mismatch.

In addition, the model provides insight into a number of further experimental observations and puts forward testable predictions:

1. the matching speed increases with initial ocular dominance, suggesting ocular dominance as a key driver of the binocular matching process.
2. matching proceeds faster than the sharpening of the orientation selectivity, suggesting that matching is not driven by the orientation selectivity.
3. the matching speed decreases with greater initial orientation selectivity.
4. the orientation selectivity becomes enhanced through the matching process only when the mismatch is sufficiently small.
5. there are two main routes to matching: the preferred orientations either drift towards each other or one of the orientations switches quite suddenly, involving a transient loss of binocularity, which can become permanent if it occurs towards the end of the critical period. While drifting occurs for small initial mismatch, switching is specific for large mismatch.

We expect that these results provide insight more generally into how neuronal systems can develop to integrate inputs from multiple sources coherently in order to generate normal neuronal function.

## 2 Methods

### Neuron model

We used an adaptive exponential integrate-and-fire model [13] with an additional current describing an afterpotential depolarization [14]. In this model the evolution of the postsynaptic membrane potential *u*(*t*) was given by

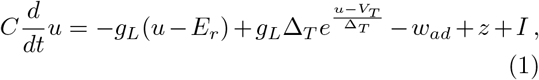

where *E*_*r*_ was approximately the resting potential, *C* the membrane capacitance, *g*_*L*_ the leak conductance, and *I* the current stimulation. The exponential term mimicked the activation of sodium current. Upon reaching the peak voltage *V*_*peak*_, the voltage *u* was reset to the fixed value *V*_*reset*_. The parameter Δ_*T*_ was the slope factor and *V*_*T*_ was the (variable) threshold potential. The variable *w*_*ad*_ represented a hyperpolarizing ptation current with dynamics given by

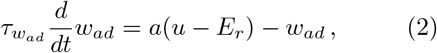

where 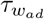 was the time constant of the adaptation of the neuron and *a* controlled the strength with which *w*_*ad*_ was driven. On firing, *w*_*ad*_ was increased by an amount *b*. The afterpotential depolarization was captured by the variable *z*. It was set to *I*_*sp*_ immediately after a spike and decayed then with a time constant *τ*_*z*_,

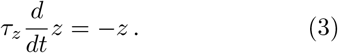

Refractoriness was modeled by employing an adaptive threshold *V*_*T*_, which was set to 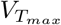 immediately after a spike and decayed then to 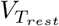 with a time constant 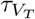,

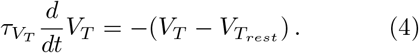

Parameters for the neuron were taken from [15] and kept fixed throughout all simulations (see Table 1).

**Table 1:** Parameters used in the model for the neuron. All parameters were set in advance on the basis of [15].

| Parameter | Value | Parameter | Value |
| --- | --- | --- | --- |
| $C$, membrane capacitance | 281 pF | $a$, subthreshold adaptation | 4 nS |
| $g_L$, leak conductance | 35 nS | $b$, spike triggered adaptation | 0.0805 nA |
| $E_r$, approximated resting potential | -70.6 mV | $I_{sp}$, spike current after a spike | 400 pA |
| $\Delta_T$, slope factor | 2 mV | $\tau_z$, spike current time constant | 40 ms |
| $V_T$, threshold potential at rest | -50.4 mV | $\tau_{V_T}$, threshold potential time constant | 50 ms |
| $V_{reset}$, resetting voltage | -50.4 mV | $V_{T_{max}}$, threshold potential after a spike | 30.4 mV |
| $V_{peak}$, spiking threshold | 20 mV | $k$, argument of modified Bessel function for synaptic input | 1.7 |
| $\tau_{w_{ad}}$, adaptation time constant | 144 ms | $A$, amplitude of the orientation-selective response of synapses | 0.14 |
| $g_{ex}$, excitatory synaptic conductance | 35 nS | $V_{ex}$, reversal potential of the excitatory synapse | 0 mV |
| $g_{inh}$, inhibitory synaptic conductance | 40 nS | $V_{inh}$, reversal potential of the inhibitory synapse | -80 mV |

To test the robustness of our results we also used a simplified neuron model with both the adaptation current and afterdepolarization removed, with which we obtained very similar results.

### Synaptic Inputs

Our model consisted of one postsynaptic binocular cell modeling a cortical cell in V1 receiving 500 excitatory, monocular, tuned synaptic inputs (Fig.1A), driven by independent Poisson spike trains. They were divided equally into inputs from the left and the right eye, respectively. In addition, inhibitory, untuned synaptic inputs were introduced to capture the sublinear binocular integration observed experimentally [16] (Fig.2). The monocular orientation preferences of the tuned excitatory synapses were linearly spaced between 0° and 180°. To mimic visual input consisting of gratings oriented at an angle *θ*_0_ each excitatory synapse *i* with preferred orientation *θ*_*i*_ received as input a Poisson spike train with an average firing rate given by the von Mises distribution with center 2*θ*_0_,

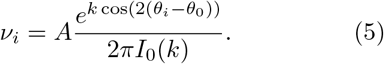

**Figure 1:**
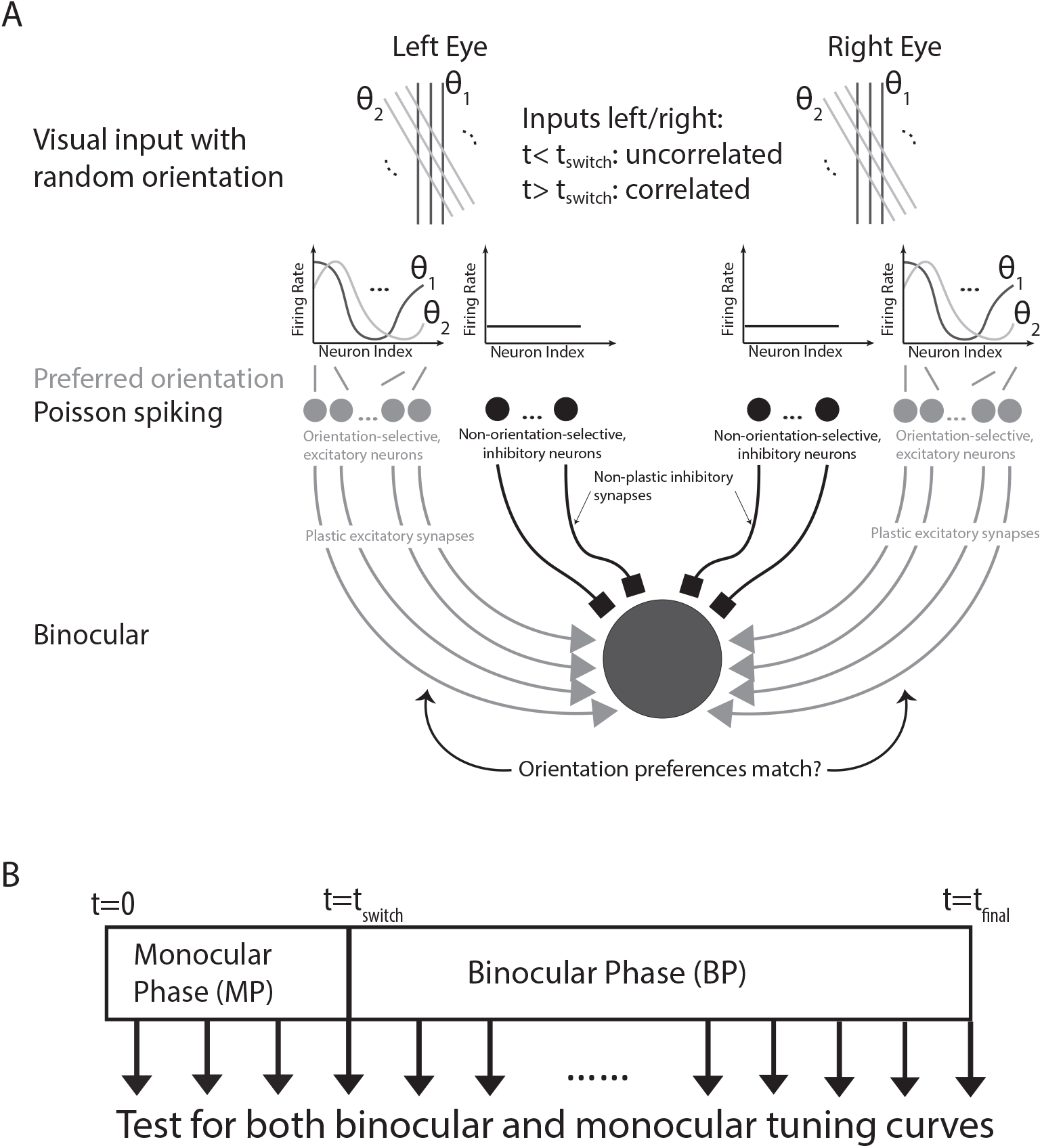
Computational Model. (A) The postsynaptic neuron received synaptic inputs as Poisson spike trains from 500 excitatory, orientation-selective synapses and conductance-based current inputs from non-selective inhibitory synapses, divided equally into inputs from the left and the right eye. Random sequences of oriented visual input characterized by orientation *θ* were presented to each eye. Only excitatory synapses were plastic. (B) Simulation protocol: The orientation of the simulated visual input was randomly shifted every 225 ms. The left and right inputs were uncorrelated until *t*_switch_ = 56.25*s* (monocular phase), then left and right inputs were identical (binocular phase).

**Figure 2:**
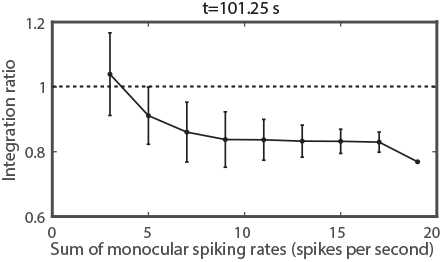
Sublinear integration ratio of the spiking rate. The ratio of the spiking rate for binocular input during the binocular phase and the sum of the corresponding monocular rates is plotted against the sum of the monocular spiking rates.

Here the modified Bessel function of order 0, *I*_0_(*k*), provided the normalization and *A* controlled the overall amplitude of the input. The value of *k* was determined by matching the tuning width of *v*_*i*_ to that observed for neurons in layer 2/3 [17, 18]. All excitatory and inhibitory synapses delivered conductance-based currents. The total synaptic current *I*_*syn*_ was given by

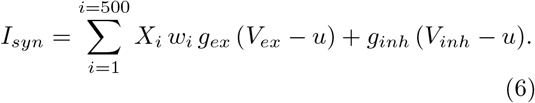

where *g*_*ex*_ (*g*_*inh*_) was the excitatory (inhibitory) synaptic conductance, *V*_*ex*_ (*V*_*inh*_) the reversal potential of the excitatory (inhibitory) synapses. The presynaptic Poisson spike trains were given by 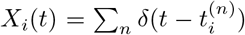 with *i* the index of the synapse and *n* counting the spikes in the train. The conductance *g*_*inh*_ of the inhibitory synapses was chosen to match the experimentally observed binocular sublinear integration ratio [16] (Fig.2). Since the inhibitory synapses were not plastic, the timing of that input was not essential and we modeled it as steady. The difference in the strength of input from the ipsilateral eye and from the contralateral eye was not included in the model.

Ocular dominance has been revealed as playing a key role in the matching outcome [12]. In experiments the ODIs of V1 cells that are upstream from the neuron in our model are distributed between −1 and +1 with a small bias toward the contralateral eye [19]. To focus on the key elements of the development and binocular matching of monocular preferred orientation and to avoid unnecessary complexities, we restricted in our model the ODI of the upstream neurons to the extreme values ±1. Thus, the upstream cells were effectively assumed to be monocular. Correspondingly, we did not include in our model an early phase during which the preferred orientations of upstream neurons become matched.

### Plasticity model

The excitatory synapses were chosen to be plastic while the strength of the inhibitory synapses were kept fixed. As plasticity model we chose the model of voltage-based STDP with homeostasis introduced in [15], which exhibited separate additive contributions to the plasticity rule for long-term depression (LTD) and for long-term potentiation (LTP),

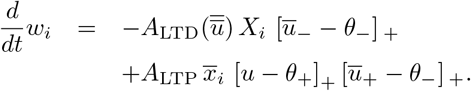

The weights were limited by hard bounds, *w*_*min*_ ≤ *w*_*i*_ ≤ *w*_*max*_. The LTP component depended on the postsynaptic membrane potential and a low-pass filtered version of the presynaptic spike train obtained via

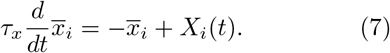

The low-pass filtered, postsynaptic membrane potentials 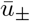 were obtained via

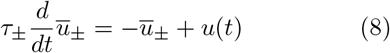

and entered the plasticity rule through the rectifier denoted by […]_+_. The amplitude 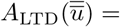 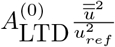 captured a homeostatic process based on the low-pass filtered square of the deviation of the membrane potential from the resting potential,

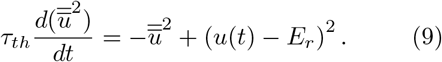

Thus, the key features of this plasticity model are that depression occurs when a presynaptic spike arrives and the average voltage 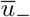 surpasses the threshold *θ*_−_, while a synapse is potentiated if the momentary postsynaptic voltage *u*(*t*) is above the threshold *θ*_+_ and the average voltage 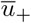 is above *θ*_−_ during a time of order *τ*_*x*_ after a presynaptic spike.

Parameters for the plasticity model were kept fixed throughout all simulations (see Table 2).

**Table 2:** Parameters of the plasticity model. Parameters as used in [15] for visual cortex, except for 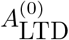, *A*_LTP_, which have been increased to speed up the simulations.

| Parameter | Value | Parameter | Value |
| --- | --- | --- | --- |
| $\theta_+$, threshold potential for voltage | -45.3 mV | $\tau_-$, time constant for filtered voltage $\bar{u}_-$ | 10 ms |
| $\theta_-$, threshold potential for filtered voltage | -70.6 mV | $\tau_+$, time constant for filtered voltage $\bar{u}_+$ | 7 ms |
| $A_{\text{LTD}}^{(0)}$, amplitude of LTD | $7 \times 10^{-4} \text{mV}^{-2}$ | $\tau_x$, time constant for the presynaptic spiking trace | 15 ms |
| $A_{\text{LTP}}$, amplitude of LTP | $12 \times 10^{-4} \text{mV}^{-2}$ | $\tau_{th}$, time constant for the homeostasis | 1.2 s |
| $w_{min}$, lower bound for the synaptic weight | 0 | $w_{max}$, upper bound for the synaptic weight | 1.6 |

### Simulation

The initial strengths of the excitatory synapses were chosen randomly from a uniform distribution within [*w*_*min*_, *w*_*max*_]. For the first stage from *t* = 0 to *t* = *t*_switch_, we simulated monocular vision by presenting a random sequence of oriented visual inputs that were uncorrelated between the left and the right eye. The orientation of the visual input was randomly changed every 225 ms. This represented the monocular phase (MP) of the simulation (Fig.1B). Then, in the second, binocular phase (BP) from *t* = *t*_switch_ to *t* = *t*_final_, we simulated binocular vision by presenting a random sequence of oriented visual inputs that were identical for the two eyes. The orientation of the visual input was again changed randomly at the same frequency as in MP. We omitted the transition period (P15-P20) during which spontaneous activity and visually evoked activity are both driving plasticity [10]. To monitor the evolution of the orientation preference we recorded all synaptic strengths every 250 ms. Monocular and binocular tuning curves were generated by testing the spiking response of the postsynaptic cell for the recorded synaptic strengths every 20s. To gather statistics, we ran the simulations multiple times (*n* = 5600 trials).

All numerical simulations were performed with MATLAB. The code is available from the authors upon request.

### Data analysis

We characterized the response of the postsynaptic neuron using the average spiking rate during windows with a duration of 1 second, both monocularly and binocularly. The tuning curve was generated by plotting the response magnitude against the orientation of the visual input. We defined the orientation preference of the cell as the orientation that gave the largest response. This was done for monocular input yielding separate preferred orientations *O*_*L,R*_ for the left and right eye, respectively, and for binocular input resulting in *O*_*bino*_. The monocular/binocular spiking rate was defined as the response for the preferred orientations *O*_*L,R*_ and *O*_*bino*_, respectively. The mismatch Δ*O* in the orientation preference was calculated as the smaller of the two values |*O*_*L*_ − *O*_*R*_| and 180° − |*O*_*L*_ − *O*_*R*_|. The global orientation selectivity index (gOSI) was computed as the magnitude of the sum 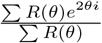 over all angles with *R*(*θ*) giving the firing rate response at orientation *θ*. The ocular dominance index (ODI) for each cell was calculated as 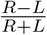, where *R* and *L* represent the maximum response magnitude for input from the right and left eye, respectively. The ODI ranges from −1 and 1, where positive values indicate right bias and negative values indicate left bias. The prediction error for the matching outcome was the difference between the predicted and the measured binocular preferred orientation. The decay rate of mismatch during binocular vision from *t*_1_ (with mismatch Δ*O*_1_) to *t*_2_ (with mismatch Δ*O*_2_) was determined by 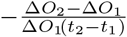. To reduce the impact of noise, we included in the computation of the matching rate only times for which Δ*O*_1_ was larger than 30°.

## 3 Results

Here we developed a simple computational model (Fig.1A) to investigate the development and matching of receptive fields in a multi-channel system in the context of the orientation selectivity and its binocular matching in primary visual cortex (V1). The model consisted of one hypothetical binocular cell receiving 500 excitatory, monocularly tuned synaptic inputs, which were divided equally into inputs from the left and the right eye. The orientation preferences of the inputs from each eye were linearly spaced in the range 0° to 180°. Thus, each synapse received an independent Poisson spike train with a firing rate that depended on the difference between its preferred orientation and the orientation of the simulated input. In addition, the post-synaptic cell received inhibitory, untuned synaptic inputs to capture the experimentally observed sublinear binocular integration [16] (Fig.2). These inputs were also modeled as Poisson spike trains. While the inhibitory synapses were non-plastic, the excitatory synapses were plastic; their synaptic weights changed depending on the timing of pre- and post-synaptic spikes, the post-synaptic voltage and low-pass filtered versions thereof [15]. Note that, effectively, the visual inputs in our model were all presented with the same spatial phase. Therefore, the difference in the phase dependence of the response of complex cells and simple cells in V1 was not considered in the model.

We started the computation with uniformly distributed random synaptic weights. In the first, monocular phase (MP) from time *t* = 0 to *t* = *t*_switch_ we simulated monocular vision by presenting inputs that corresponded to bars with an orientation that was uncorrelated between the left and the right eye and randomly switched every 225ms. In the second, binocular phase (BP) from *t* = *t*_switch_ to *t* = *t*_final_ we simulated binocular vision by presenting identical visual inputs to the two eyes, again randomly switching orientation in time with the same frequency as in MP. To monitor the evolution of the orientation preference of the cell, we recorded all synaptic strengths every 250ms and generated tuning curves by measuring the spiking response of the post-synaptic neuron during both MP and BP (Fig.1B). To gather statistics, we ran many trials, each resulting in an effectively different cell with different response properties (*n* = 5600 trials).

### Synaptic plasticity captures binocular matching

The results obtained in our model are consistent with key aspects of previous experiments [11, 7, 12]. Using a tuning width for the inputs that corresponds to that of cells in layer 4 [17], our model reproduced the development of orientation selectivity for V1 cells with global orientation selectivity index (gOSI) and tuning width similar to those found experimentally in layer 2/3 [17, 18]. Also, the experimentally observed sublinear binocular integration [16] was captured in our model (Fig.2).

Moreover, while right after eye-opening a fraction of V1 cells has been observed to have well-developed orientation selectivity, their monocular preferred orientations for input from the left and the right were poorly matched [11]. In fact, in some cells, they were nearly 90° apart, the maximal possible difference. This mismatch decreased substantially with age to reach the adult level by P30-P36 [11, 7].

In our model, during the initial phase of MP multiple sets of synapses were potentiated. Due to the random distribution of the initial synaptic strengths, the randomly chosen orientations of the first several inputs, as well as the variability of the number of spikes in the Poisson spike trains received by the synapses during the presentations of each input their strengths did not vary smoothly with orientation. Nevertheless, the sets of potentiated synapses roughly specified monoc ular orientational receptive fields (ORFs) of the postsynaptic neuron, defined as those orientations to which the neuron responded significantly. During MP these ORFs for input from the left and the right eye did not match (Fig.3A1,2 up to *t* = *t*_switch_, marked by white dashed lines), which manifested itself also in non-matching orientation tuning curves (Fig.3B1,2).

**Figure 3:**
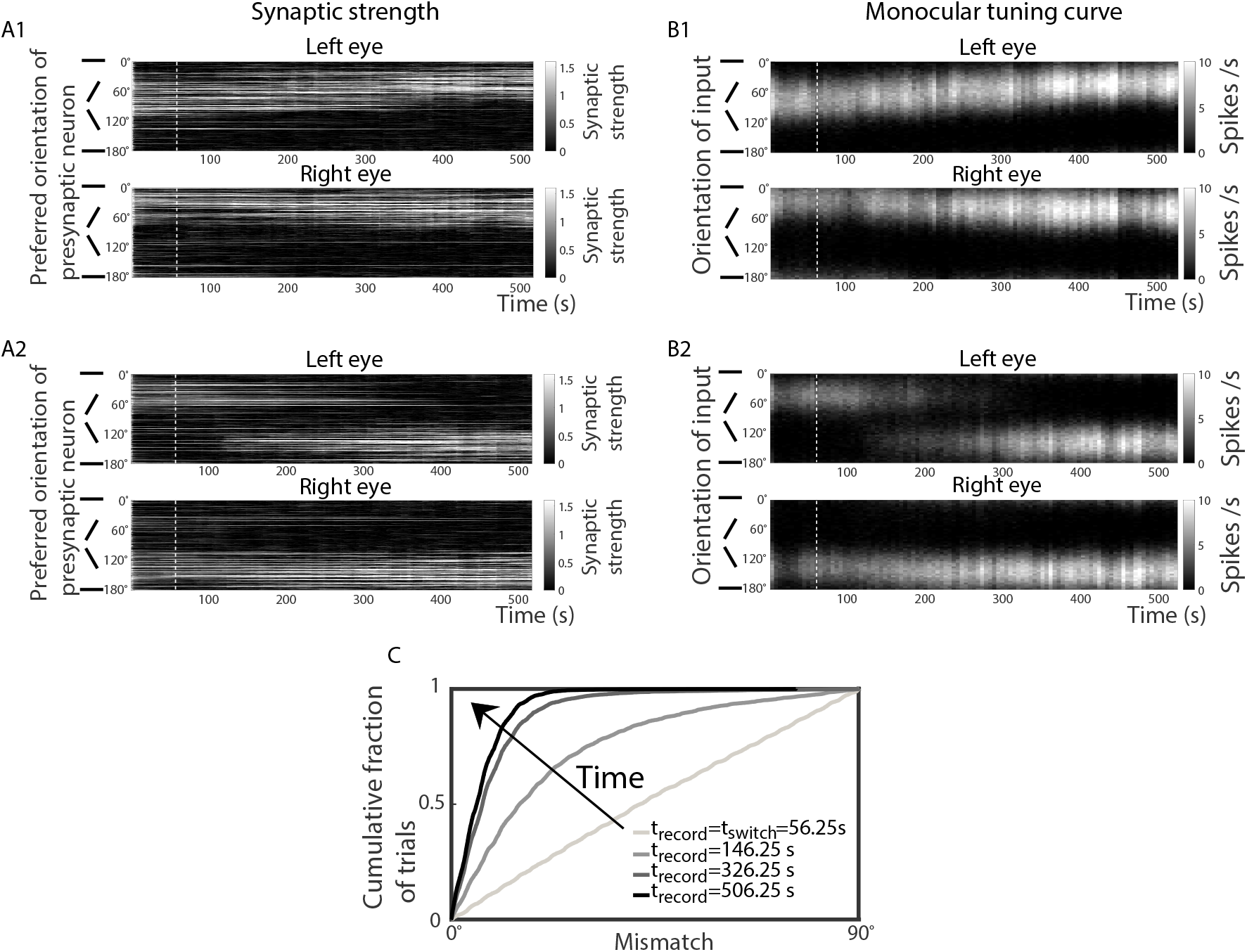
Binocular vision reduces the mismatch of the monocular orientation preferences. (A1,A2): Two examples of the temporal evolution of the synaptic strengths. The presynaptic neurons are ordered according to their preferred orientation (vertical axis), and their synaptic strengths are illustrated in greyscale as a function of time (horizontal axis). (B1,B2): Corresponding evolution of the monocular tuning curves through each eye. Greyscale indicates the firing rate of the cell in response to monocular input with an orientation as indicated along the vertical axis. In (A1,B1) the two monocular orientation preferences drifted toward each other while in (A2,B2) one monocular orientation preference switched to the other discontinuously to achieve binocular matching. White dashed lines mark *t* = *t*_switch_. (C): The cumulative distribution function of the mismatch (*t*_switch_ = 56.25s; *n* = 5600 trials).

During BP both eyes received the same inputs. This allowed the potentiation of weak synapses that by themselves were not strong enough to drive a post-synaptic spike, if the synapses with the same orientation preference but receiving input from the other eye were sufficiently strong to trigger a spike. This slowly modified the ORFs and the tuning curves (Fig.3A,B for *t* > *t*_*switch*_), decreasing the mismatch between the two monocular orientation preferences. Eventually, in almost all trials the preferences became matched within 20° (Fig.3C).

### The effect of ODI on the matching outcome

It has been shown that three weeks of environmental enrichment (EE) can rescue the disrupted binocular mismatch caused by visual deprivation during the critical period [12]. These experiments revealed that ocular dominance plays a key role in the binocular matching process. In cells whose response was dominated by one of the two eyes, binocular matching was achieved by the orientation preference for input from the nondominant eye changing, while the orientation preference for input from the dominant eye did not change much.

Motivated by this experimental result, we determined for each trial the change *δO*_*L,R*_ in the left and right monocular orientation preferences during BP as well as the ocular dominance index (ODI) right before BP. The statistics of *δO*_*L,R*_ and ODI across many trials are shown in two-dimensional histograms (Fig.4A,B). As in the experiments, the range in the change of orientation preference for input from the nondominant eye was much wider than that for the dominant eye. Fig.4C shows an illustrative example in which at *t*_*switch*_ (white dashed line) the cell was dominated by the input from the right eye (ODI= 0.415). During BP the preferred orientation for input from the nondominant (left) eye changed substantially, while that for input from the dominant (right) eye did not evolve much.

**Figure 4:**
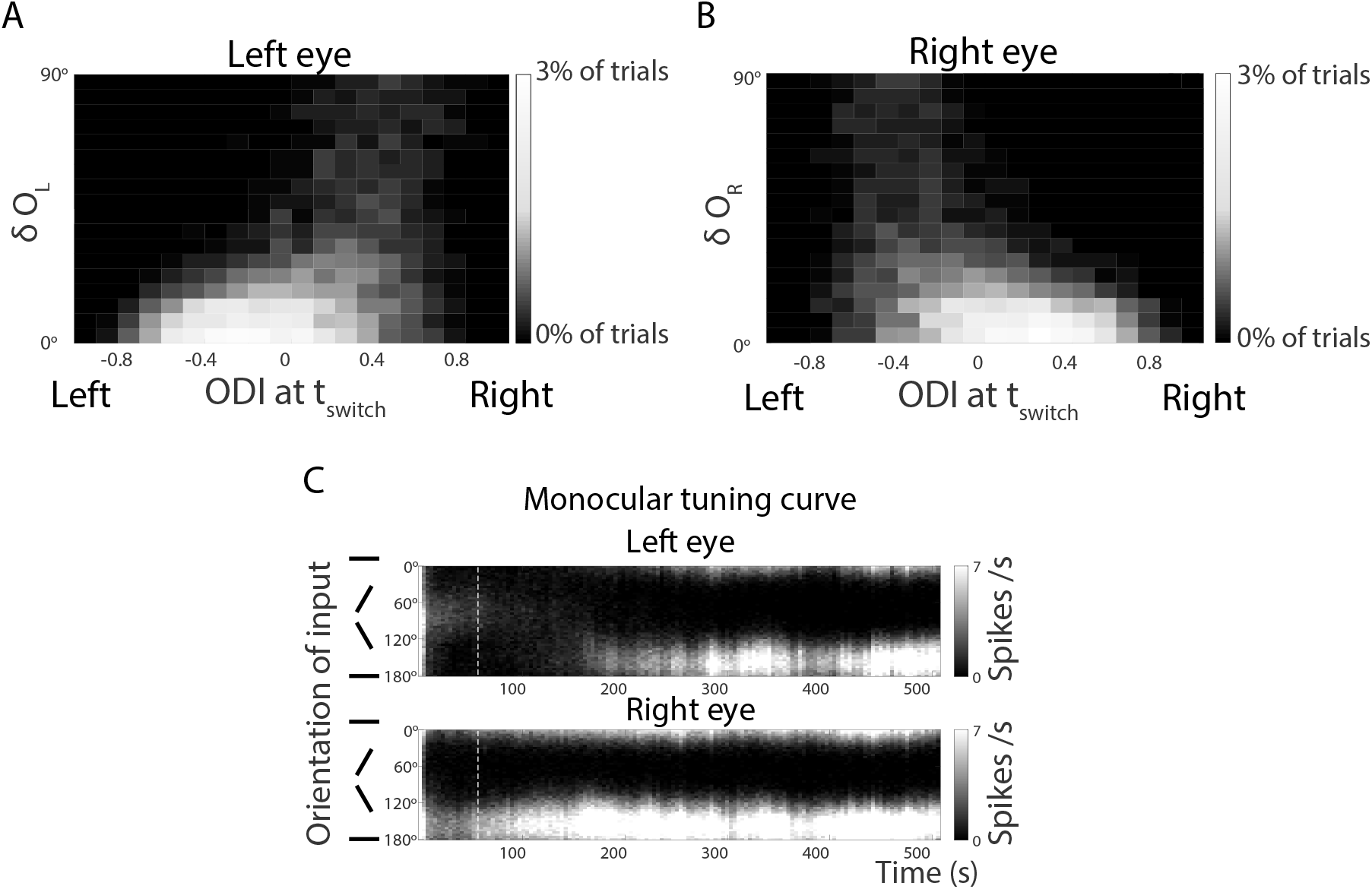
Orientation preference mostly changes for nondominant eye. (A) Two-dimensional histogram in which the greyscale of each square bin indicates the percentage of trials yielding cells whose monocular orientation preference through the left eye changed by *δO*_*L*_ (vertical axis) during BP and that had an ODI at *t*_*switch*_ as given on the horizontal axis. (B) Same plot as (A), except for the right eye. In both (A) and (B) *t*_switch_ = 56.25s, *t*_final_ = 506.25s, *n* = 5600 trials. (C) The evolution of the monocular tuning curve for input from the left and right eye (cf. Fig.3B). White dashed lines mark *t* = *t*_switch_. The scale of the greyscale map is capped to better show the difference between the left and right monocular firing rates at *t*_*switch*_ (ODI>0).

### The interaction between orientation selectivity and matching

Previous experimental results revealed an inverse relationship between mismatch and gOSI: cells with smaller orientation mismatch had greater orientation selectivity. This did not hold in mice whose binocular matching process was compromised by visual deprivation; their Δ*O* values spanned the entire 0°–90° range for all gOSI values [12]. The histograms in Fig.5 show the relationship between mismatch and gOSI obtained in the model at the onset of BP at *t*_*switch*_ (Fig.5A) and at an intermediate time during BP (Fig.5B). In most trials, the cell was neither well-matched nor very selective at the end of MP (Fig.5A). Consistent with the experimental results, at intermediate times during BP the mismatch was small in highly orientation-selective cells (Fig.5B).

**Figure 5:**
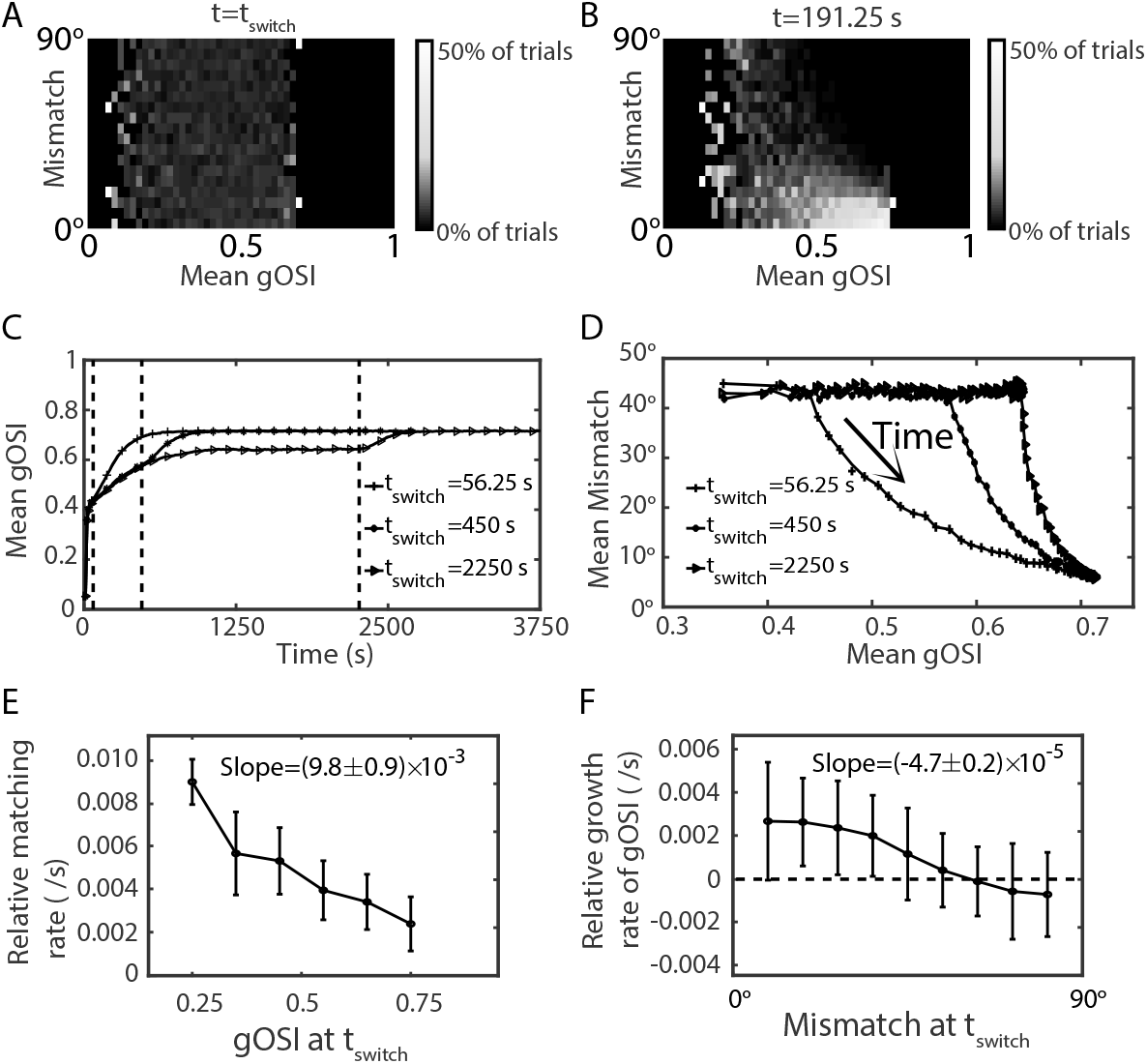
Matching leads the development of orientation selectivity during BP. (A,B): Histogram of the mismatch for different values of the mean monocular gOSI at *t* = *t*_switch_ = 56.25s. The histogram is normalized for each value of the mean monocular gOSI. (B) As (A), but for *t* = 191.25s during BP. Lower mismatch is correlated with higher orientation selectivity (*n* = 5600 trials). (C) Binocular vision enhances orientation selectivity. The evolution of gOSI for *t*_*switch*_ = 56.25s, 450s, 2250s (marked by dashed lines). (*n* = 1000 trials). (D) Mismatch vs. gOSI during the evolution shown in (C). Orientation selectivity lags behind the matching during BP (*n* = 1000 trials, *t*_*final*_ = *t*_*switch*_ + 562.5*s*). (E) Less selective cells match faster. The relative matching rate is given by 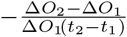, where Δ*O*_1,2_ is the mismatch at time *t*_1_ = *t*_*switch*_ = 56.25s and *t*_2_ = 101.25s, respectively. (F) Cells do not become more selective unless the mismatch is small enough. The relative growth rate of gOSI is given by 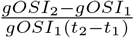 where *gOSI*_1,2_ is measured at time *t*_1_ = *t*_*switch*_ = 56.25s and *t*_2_ = 101.25s, respectively. In panel E and F the legend gives the slope of a linear regression of the data to quantify the trend.

To gain insight into the relationship between the matching process and the sharpening of the orientation selectivity, we measured the evolution of the gOSI for different durations of the MP. For *t*_*switch*_ > 1250s the gOSI reached a steady state during MP (Fig.5C). Remarkably, its saturation value was significantly lower than the value reached during BP, even if that BP followed an MP with short duration. This indicates that binocular vision enhanced the development of orientation selectivity. Moreover, the mismatch approached its final value faster than the gOSI did (Fig.5D). Further analysis employing initial conditions with well-controlled mismatch showed that less selective cells matched faster (Fig.5E) and cells did not become more selective until the mismatch had become small enough (Fig.5F). This suggests that the matching process enhanced the orientation selectivity, while orientation selectivity was not a driving force of binocular matching but had, instead, a negative effect on matching.

Moreover, at the onset of BP the left and right orientation selectivities were often quite different from each other leading to a broad distribution across trials (Fig.6A). But binocular vision enhanced the selectivities and drove them to the same large value (Fig.6B,C).

**Figure 6:**
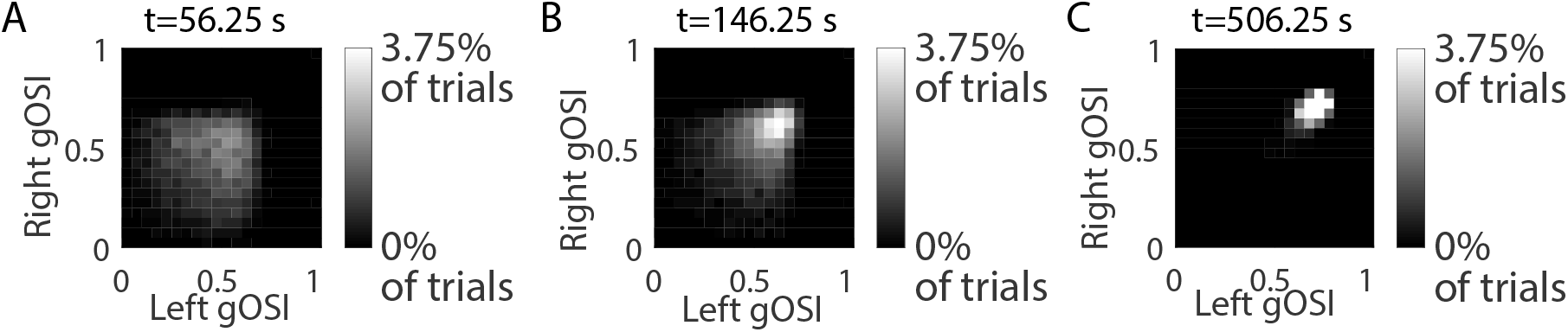
Binocular vision enhances and matches monocular orientation selectivity. Two-dimensional histograms of right and left gOSI at the onset (A, *t*_*switch*_ = 56.25s) and two intermediate times during BP (B, C). (*n* = 5600 trials)

### Prediction of matching outcome

Next, we put forward testable predictions based on our model. We hypothesized that the binocular orientation preference right before BP could predict the eventual matching outcome. We therefore defined the prediction error as the difference between the final binocular preferred orientation and that at the beginning of BP (Fig.7A). Overall, the binocular orientation preference at *t*_*switch*_ was a quite good predictor for the orientation preference at all time points during BP that we investigated (Fig.7B). Note that in Fig.7B the histogram of the prediction error was normalized separately for each value of the mismatch. This revealed that for small mismatch almost all trials had a prediction error of less than 20° while for large mismatch the distribution of prediction errors was almost uniform. Thus, the mismatch that remained at a given time during BP indicated quite well whether the binocular orientation preference at *t*_*switch*_ was a good predictor for the binocular orientation preference at that later time.

**Figure 7:**
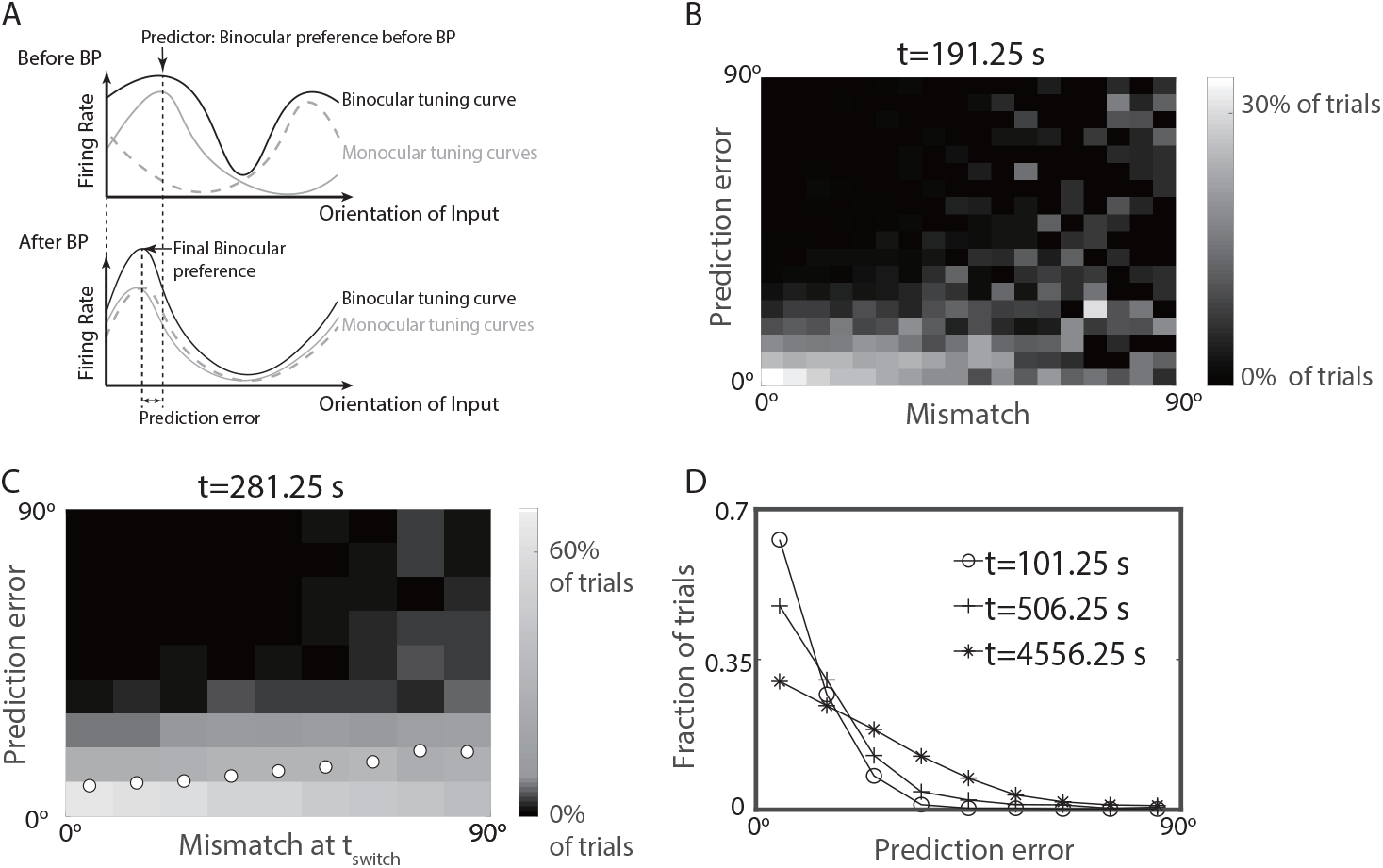
Binocular orientation preference at the onset of BP predicts the preference after matching. (A): The prediction error was calculated as the difference between the binocular orientation preference before and after BP. (B): The quality of the prediction decreased with increasing mismatch. Histogram of the prediction error for different values of the mismatch, both at the same time *t* = 191.25s. The histogram is normalized separately for each value of the mismatch. (C): The accuracy of prediction decreased with the *initial* mismatch. Like (B), but at *t* = 281.25s and using the *initial* mismatch. Circles represent the mean prediction error of the trials for each value of initial mismatch. (D): The mean prediction error increased with time. The distribution of the prediction error from well-matched trials (mismatch < 20°) at three intermediate times during BP. In all figures, *t*_switch_ = 56.25s and *n* = 5600 trials.

Can the reliability of the prediction for the preferred orientation already be anticipated at *t*_*switch*_? Indeed, already the *initial* mismatch was a good indicator for this reliability (Fig.7C): the prediction was quite accurate when the initial mismatch was small, while it became less reliable for large initial mismatch.

Since in the experiments the monocular, rather than the binocular tuning curves were measured [12], we tested how well the superposition of two monocular tuning curves at the onset of BP could predict the final, matched orientation. We found that at each time point the orientation preference determined by the linear superposition of two monocular curves was close to the binocular orientation preference and therefore it was also a good predictor for the matching outcome (Fig.8).

**Figure 8:**
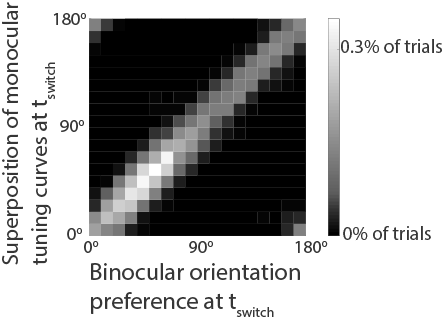
Binocular tuning is well approximated by the sum of the monocular tuning curves. Histogram of the binocular preferred orientation and the preferred orientation obtained from the sum of the two monocular tuning curves.

The matched preferred orientation that emerges in a given trial reflects the initial synaptic weights as well as the sequence of presented visual inputs. Since in our simulations the synapses were plastic throughout the simulation and the sequence of inputs random, the binocular preferred orientation evolved on a slow time scale, wandering around in a diffusive manner. The distribution of the prediction error across trials therefore broadened with time as shown in Fig.7D for well-matched cells (mismatch less than 20°), implying a growing mean prediction error.

### The speed of matching depends on the initial ODI

We next examined how initial ocular dominance affected the speed of the matching process. Figs.9A1,2,3 show the histogram of the mismatch for various time points as a function of the magnitude |ODI| of the ocular dominance index at *t*_*switch*_. For high initial |ODI| the mismatch decreased rapidly, while in many trials that had a lower initial |ODI| the matching proceeded more slowly (most clearly seen comparing *t* = 146.25s with *t* = *t*_*switch*_). We quantified this in terms of the decay rate of the mismatch given by 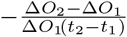, where Δ*O*_1,2_ is the mismatch at time *t*_1,2_, respectively. Fig.9B shows the mean and standard deviation of the decay rates for different ranges of the |ODI| at *t*_*switch*_. This result is consistent with the intuition that cells with a low initial |ODI| have two monocular ORFs with similar overall synaptic strengths, which compete with each other during BP, slowing down the matching process. This effect of ocular dominance on the binocular matching rate reveals ocular dominance as a driver of the binocular matching process. Note that trials with an initial mismatch less than 40° were not included in Fig.9B, since here we were only interested in the matching processes starting with a state that was not well-matched. Similar values for the decay rates as shown in Fig.9B were obtained at other intermediate times during BP, suggesting an exponential decay of mismatch during BP.

**Figure 9:**
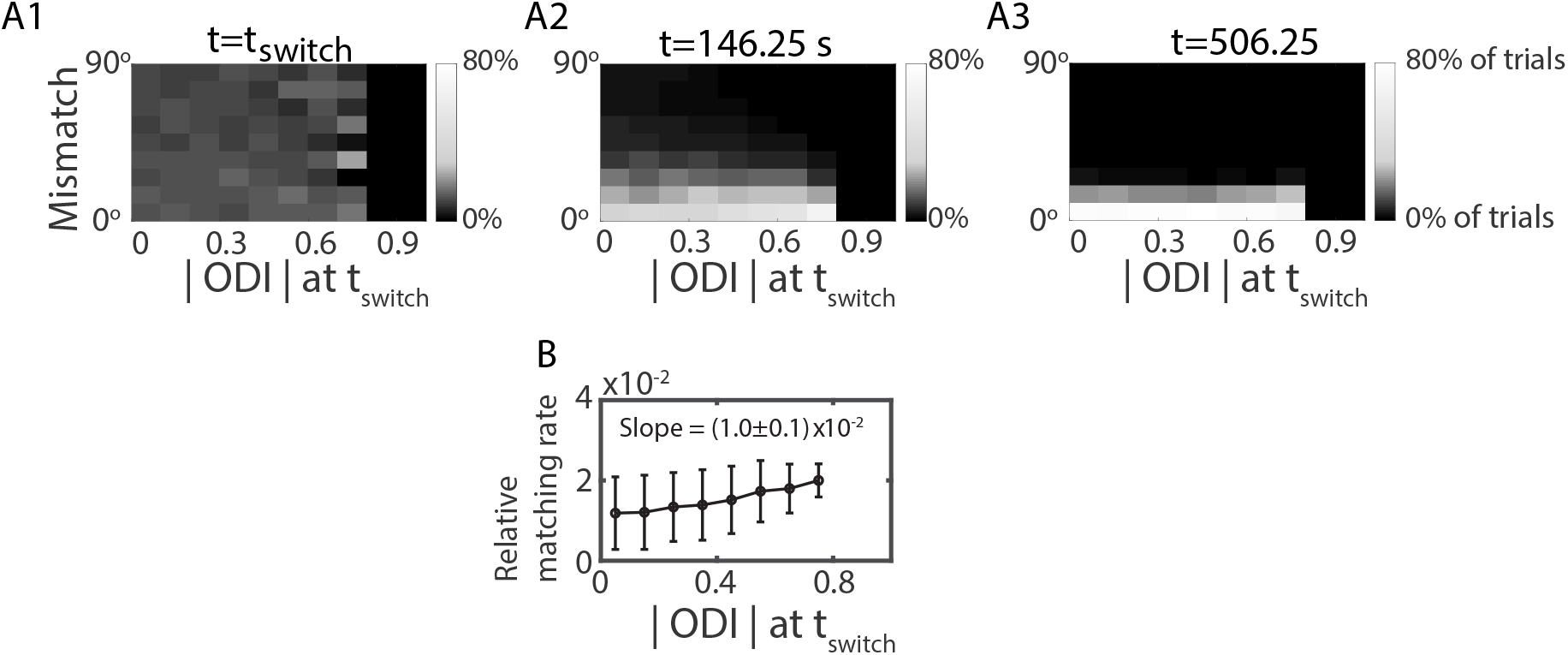
Binocular matching speed increases with the absolute value of initial ODI. (A1,2,3): Histograms of mismatch at different times for different values of the magnitude of ODI at *t*_*switch*_ (*t*_switch_ = 56.25s, *n* = 5600 trials). The histograms are normalized for each value of initial |ODI| separately. (B): The average relative matching rate during the time interval [56.25s, 146.25s] against the ODI at *t*_switch_ = 56.25s (3537 out of 5600 trials). The legend gives the slope of a linear regression of the data to quantify the trend.

### The type of matching process depends on the initial mismatch

Finally, we investigated whether there are qualitatively different processes through which the binocular matching was obtained. Figs.3B1,2 show that there are at least two ways to achieve binocular matching. In Fig.3B1, the two monocular orientation preferences drifted towards each other, whereas in Fig.3B2 one orientation preference switched to the other one discontinuously. To determine whether the matching process is achieved by drifting or switching, we examined the temporal evolution of the monocular gOSI of each eye during BP. When the two monocular orientation preferences drifted towards each other, the cell was orientation-selective for input from each eye throughout the matching process, mean ing that both monocular gOSIs stayed high at all times (Fig.10A). In contrast, when the matching was achieved by switching, the monocular ORF for input from one eye disappeared at some time and a new one gradually appeared, which matched the monocular ORF for input from the other eye. In this case, there was a period during which the cell was not orientation-selective for input from one eye, i.e. one monocular gOSI was low (Fig.10B).

**Figure 10:**
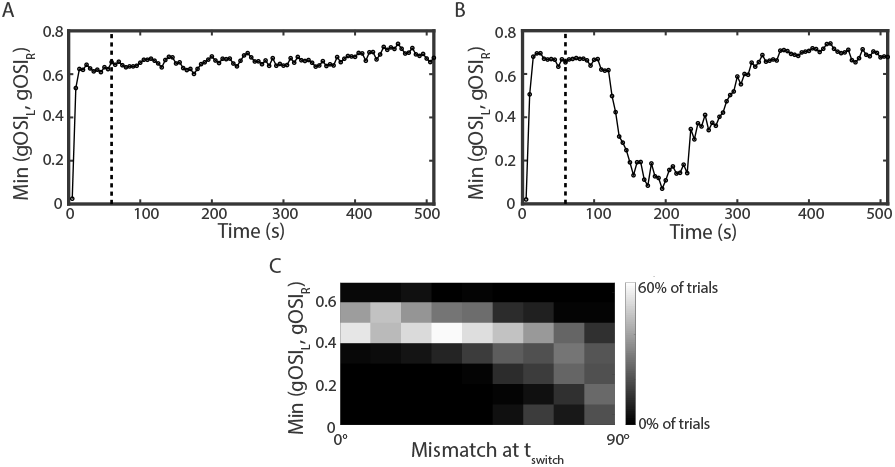
The type of matching depends on the initial mismatch. (A) The evolution of the smaller of the left and right monocular gOSIs for Fig.3B1. Both gOSIs remained high throughout BP. (B) Like (A) for Fig.3B2. One of the gOSIs dropped transiently during BP. In both (A) and (B), dashed lines mark *t* = *t*_*switch*_. (C) Histogram of the minimal value of the monocular gOSIs across the duration of BP for different values of the mismatch at the onset of BP. Only trials with monocular gOSIs through each eye higher than 0.45 at the onset of BP were included (1473 out of 5600 trials). The histogram was normalized for each value of the initial mismatch. The gOSIs were tested every 50s from *t*_switch_ = 562.5s to *t*_final_ = 5062.5s.

To discriminate between the two matching processes we determined the minimal monocular gOSI across the whole BP and related it to the mismatch at *t*_*switch*_ (Fig.10C). For small initial mismatch both monocular gOSIs remained large throughout the matching process, implying that binocular matching was achieved by the monocular orientation preferences drifting towards each other. If the initial mismatch was large, however, it was more likely that one of the two orientation preferences switched to the other.

Interestingly, the substantial drop in the gOSI during the switching process implied that the cell transiently responded only weakly to the input from one eye; it became effectively monocular during that transient.

## 4 Discussion

Using computational modeling we have investigated the development of orientation selectivity and its binocular matching in V1. The model focused on a hypothetical neuron that receives inputs from both eyes via plastic synapses. Motivated by the fact that in a multi-source system upstream neurons gain selectivity before the downstream neurons [7, 1], we chose the inputs from the upstream neurons to be orientation selective. For simplicity we took the upstream cells to be monocular. The setup of the model cell is reminiscent, but not identical, to that of a V1 complex cell receiving oriented inputs from simple cells. Experimentally, simple cells are typically binocular and become matched earlier during the critical period (P22-P23) than complex cells (P26-P31) [7]. Thus, most inputs that complex cells receive from individual simple cells are already well matched during the relevant time period. However, simple cells vary substantially in their ocular dominance. Thus, simple cells with different preferred orientations are likely to be dominated by different eyes. Effectively, these cells provide mismatched input to the complex cells. Our model corresponds then to the simplifying, extreme case in which the ODI of all presynaptic cells is either +1 or −1.

One goal of our modeling was to investigate what experimental findings can be captured parsimoniously in a minimal computational model. We therefore assumed that the synaptic plasticity mechanism itself does not change with eye-opening and the subsequent onset of the critical period for ocular dominance; only the input to the cortical neuron, which drives the synaptic plasticity, was taken to change from being uncorrelated between the two eyes before eye-opening to being correlated after eye-opening. This simplification is consistent with a number of experimental findings. In cats, orientation selectivity emerges already before eye opening, driven by vision-independent spontaneous activity in the retina [20], and continues to increase after eye opening. Until the onset of the critical period this increase does not depend on visual input [21]. Correspondingly, the onset of the critical period has been identified not as a change in the plasticity mechanism but as a transition from synaptic plasticity being driven predominantly by spontaneous activity to being driven mostly by visually evoked input [10]. This change results from an increase in inhibition, which reduces the weaker, spontaneous activity but not the stronger, visually evoked activities - to magnitudes that are not sufficient to drive synaptic plasticity. Note that this scenario may vary across species [22, 20].

Despite its simplicity, our model captured a number of fundamental experimental results for the development of orientation tuning in mouse V1 [11, 12, 7]. During the monocular phase a fraction of the cells became orientation-selective with respect to inputs from both eyes, but the preferred orientations were rarely matched. The matching occurred during the binocular phase and depended strongly on the ocular dominance of the neuron: the final binocular orientation preference was more likely to be aligned with the monocular preference for input from the initially dominant eye [12, 23]. As in the experiment [12], the orientation selectivity was higher in well-matched cells and enhanced by correlated visual input [18]. Both results reflect the enhanced drive the cells receive once the monocular tuning curves overlap, which leads to stronger plastic changes as well as enhanced selectivity due to the synaptic homeostasis [15].

Detailed analysis of the model and its results provided deeper insight into the behavior of this system and shed light on a number of further experimental observations. In the model the development of orientation selectivity and its matching are driven by different types of correlations in the inputs to the cells and therefore differ in their time course. Orientation selectivity starts to emerge already before eye opening. After eye opening visual input is expected to enhance cortical activity and with it the learning speed for orientation selectivity, consistent with the results of [18]. In the model the matching process requires correlated inputs from both eyes. It therefore suggests that matching does not start until eye-opening and persists then throughout the critical period. Indeed, experimental results suggest that at P15-P18, i.e., shortly after eye-opening, the matching of V1 cells is still close to chance level [7], which is consistent with the lack of matching in dark-reared mice [11]. However, already at the beginning of the critical period (P19-P21) the orientation preferences are somewhat matched [7]. Due to the presence of uncorrelated spontaneous activity during the phase between eye-opening and the critical period [24], the model suggests that during that phase the matching proceeds more slowly than during the critical period.

In experiments the preferred orientation is typically characterized by the orientation evoking the maximal response. Since it is predominantly the synaptic weights for inputs from the subdominant eye that change in the matching process, the preferred orientation obtained after matching is predicted to be close to the initial binocularly measured preferred orientation.

For large mismatch the matching speed is predicted to depend significantly on the ocular dominance. In the model, cells whose response was dominated by the input from one eye matched faster than cells that were equally responsive to inputs from both eyes. This reflects a competition between the inputs from the two eyes.

In the model, cells whose left and right monocular tuning curves overlap match more rapidly. Thus, for a given mismatch less selective cells are predicted to match faster. This is consistent with results obtained in mice that were reared in the dark from P1 to P30 [11]. At P30 the distribution of their mismatch was not statistically different from a uniform distribution and their selectivity was lower than that observed at the beginning of the critical period. As found in the model, their matching progressed faster than was the case for undeprived animals during the critical period. Conversely, binocular deprivation between eye-opening and the onset of the critical period has been found to increase the fraction of cells that have strong orientation selectivity [21] but large mismatch [11]. Our model predicts that their matching process will be slower.

Conversely, the model predicts that the mismatch affects the orientation selectivity. By manipulating the initial mismatch at fixed orientation selectivity, we showed that during the binocular matching process cells did not become more orientation-selective unless the mismatch was small enough to allow the monocular tuning curves to overlap.

Moreover, the overlap of the tuning curves is predicted to affect the matching process in a qualitative manner. For small mismatch, for which the tuning curves overlap significantly, the monocular preferred orientations are predicted to shift smoothly towards each other. For large mismatch, however, the model predicts that the response to input from one eye and its selectivity drop substantially during the evolution. If the plasticity period continues sufficiently long beyond that phase, this reduction in response is only transient and the response eventually recovers with a preferred orientation that has switched to that of the input from the other eye. This switching process is predicted to be more likely after binocular deprivation between eye-opening and the onset of the critical period. If the switching occurs, however, late in the critical period, the remaining duration of the plastic period may not suffice for the recovery and the cell may remain essentially monocular. This has been reported experimentally for a fraction of L2/3 cells [23].

When the plasticity period in the model was sufficiently long, all cells became highly selective and very well matched, more so than observed experimentally [11]. This could result from an over-simplification of the plasticity mechanism or of the stimuli used in the simulations. Alternatively, it could suggest that biologically the overall plasticity process and its duration are not optimized specifically for orientation selectivity and matching, but could have additional objectives. This interpretation is supported by the observation that in the model the best orientation selectivity and matching would be achieved in the shortest time if there was no monocular period at all. However, it has been pointed out that such a monocular period during which contra- and ipsilateral inputs are uncorrelated is necessary to form retinogeniculate and geniculocortical connections with segregated eye-specific areas in LGN (reviewed in [25]).

On long time scales the matched preferred orientations are not stable in the model. Instead, they perform diffusive motion, reflecting the stochastic nature of the presented inputs. To what extent this drift is biologically relevant depends on the duration of the period during which the relevant synapses are plastic. The somewhat limited degree of selectivity and matching that is observed by the end of the critical period suggests that this duration may not be long enough to observe this drift [26].

In our model the plasticity resulted from changes in the synaptic weights that were driven by correlations between presynaptic spikes and the evolution of the postsynaptic voltage, combined with a homeostatic mechanism based on the postsynaptic long-term activity [15]. We expect that most of our results do not depend on the specific details of the plasticity mechanism as long as it has a Hebbian component that is based on the correlations between pre− and postsynaptic activities and that does not change the weights for low presynaptic activity, in combination with homeostatic regulation. Conceivably, the plasticity mechanism could have a strong structural component [27], which may be quite likely at this developmental stage of the animal.

We have considered in our model only a single neuron and its feedforward inputs. In the mammalian V1 the L2/3 and the L4 neurons are, however, part of a recurrent network. In higher mammals, anatomically close V1 neurons have similar preferred orientations, giving rise to orientation maps [28, 29] and neurons with similar preferred orientations have a higher probability to be connected with each other [30]. In contrast, in mouse the spatial clustering of the preferred orientations is weaker and neurons with different preferred orientations are intermixed. The development of orientation selectivity in such a recurrently connected network of neurons has been studied extensively [31, 22, 32]. How the network affects the matching process under correlated input is, however, still an open question.

The framework of our model can readily be applied to neurons in other multi-channel systems such as binaural auditory neurons or multisensory neurons to capture the development and matching of multiple, single-channel receptive fields that represent corresponding physical properties (e.g. orientation, position). For multisensory neurons in superior colliculus, for example, it has been shown that the selectivity of multisensory neurons develops after the development of selectivity of its upstream unisensory neurons [1], which is similar to the setup in our model. The binocular vision and its ensuing matching of orientation selectivity through binocular vision in our visual cortex model corresponds to sensing the same event through different modalities simultaneously and the matching of their corresponding receptive fields. Thus, the ideas and results developed here may readily carry over to explain experimental results for the development and matching of receptive fields in other sensory cortices integrating inputs across modalities [1].

To conclude, by modeling the development and binocular matching for a hypothetical cell in visual cortex V1, we captured a host of experimental results in mouse and give several predictions. Key elements of the model are the evolution and competition of two monocular receptive fields in the presence of correlated inputs. The simplicity of this framework makes it a good candidate to investigate the interaction between selectivity, channel-dominance, and mismatch of a specific physical property at the single neuron level during the matching process in multi-source experience-dependent sensory systems.

